# Homeostatic regulation of a motor circuit through temperature sensing rather than activity sensing

**DOI:** 10.1101/2025.01.10.632419

**Authors:** Delaney Cannon, Joseph M. Santin

**Affiliations:** University of Missouri-Columbia, Division of Biological Sciences, Missouri, United States of America

## Abstract

Homeostasis is a driving principle in physiology. To achieve homeostatic control of neural activity, neurons monitor their activity levels and then initiate corrective adjustments in excitability when activity strays from a set point. However, fluctuations in the brain microenvironment, such as temperature, pH, and other ions represent some of the most common perturbations to neural function in animals. Therefore, it is unclear if activity sensing is a universal strategy for different types of perturbations or if stability may arise by sensing specific environmental cues. Here we show the respiratory network of amphibians mounts a fast homeostatic response to restore motor function following inactivity caused by cooling over the physiological range. This response was not initiated by inactivity, but rather, by temperature. Compensation involved cold-activation of the noradrenergic system *via* mechanisms that involve inhibition of the Na^+^/K^+^ ATPase, causing β-adrenoceptor signaling that enhanced network excitability. Thus, acute cooling initiates a modulatory response that opposes inactivity and enhances network excitability. As the nervous system of all animals is subjected to changes in the microenvironment, some circuits may have selected regulatory systems tuned to environmental variables in place of, or in addition to, activity-dependent control mechanisms.

**Graphical Abstract:** 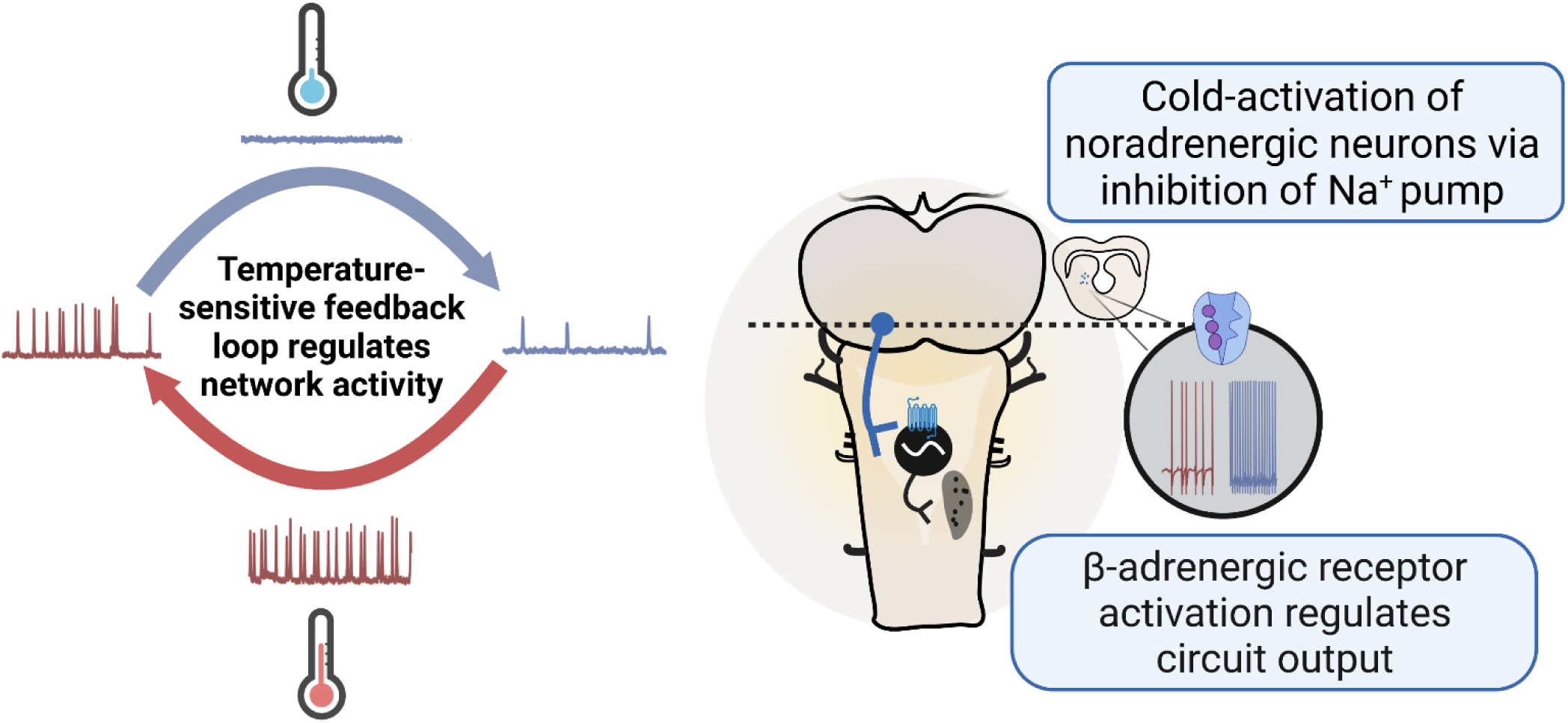

## Introduction

Negative feedback homeostasis is a core principle in physiology that explains how organisms maintain balance despite ongoing challenges in the internal and external environment (Cannon, 1929). While feedback systems regulate critical body variables like blood pH, body temperature, and hormone production, neural circuits also control activity levels through homeostasis (Wen and Turrigiano, 2024). Within this framework, neurons are thought to sense their own activity levels *via* cellular reflections of network activity, such as changes in firing rate, intracellular Ca^2+^, or neurotransmitter receptor activation (Fong et al., 2015; Ibata et al., 2008; Joseph and Turrigiano, 2017; Orr et al., 2017). During perturbations in activity, alterations in these processes serve as transduction steps that engage intracellular signaling pathways to adjust neuronal excitability in the opposite direction of the disturbance. Homeostatic processes are crucial to maintaining stable activity levels, playing a role in visual system development (Desai et al., 2002), sleep (Diering et al., 2017), emergence from hibernation (Santin et al., 2017), and formation of memory specificity (Wu et al., 2021).

Many animals inhabit environments with challenging abiotic stressors, such as fluctuations in temperature, pH, and salinity that can cause disturbances in neural function (Marder and Rue, 2021). In addition, mammals thought to have constant internal conditions also experience pronounced changes in the brain microenvironment (Chesler, 2003; Kiyatkin, 2011; Raimondo et al., 2015). While these challenges have the potential to disrupt neuronal activity, most animals persist unscathed, at least within limits. Homeostatic regulation is a logical candidate for stability during these challenges, and studies often place these processes in the context of activity-dependent signaling (Alonso et al., 2023; Rue et al., 2022; Zubov et al., 2022). However, an underappreciated possibility is that the physical environment *per se* may directly initiate regulatory responses. In principle, temperature-sensitive ion channels could produce signaling events that correct disturbances by temperature (Kashio and Tominaga, 2022; Shibasaki, 2024). The Na^+^/K^+^ ATPase and other electrogenic ion transporters have high temperature and pH dependencies and can influence network excitability through changes in the membrane potential and ion gradients (Glitsch, 2001). Finally, some G-protein coupled receptors are directly modulated by temperature, pH, and extracellular Na^+^ ions which could influence neural activity *via* metabotropic signaling (Ferré et al., 2023; Kumar et al., 2015; Ohnishi et al., 2024). Therefore, in addition to activity-dependent signals commonly implicated in feedback control, physical variables of the neuronal microenvironment may act through parallel pathways to elicit regulation.

Here, we test the hypothesis that activity-dependent signals *vs*. those from the ambient environment elicit compensation that regulates motor behavior. For this, we used the amphibian respiratory network, as these animals experience a wide range of ambient temperatures and undergo rapid changes in temperature depending on their microhabitat. To meet their metabolic demands, brainstem circuits must produce rhythmic output across a range of temperatures. However, temperatures below ∼15°C strongly depress output (Saunders and Santin, 2024; Vallejo et al., 2018), which could be catastrophic on the short term as breathing must continue for metabolic homeostasis (Gottlieb and Jackson, 1976). To overcome this challenge, we show here that inactivity caused by acute cooling induces a response consistent with homeostatic regulation over the timescale of tens of minutes to enhance excitability of the respiratory network. Rather than activity-dependent signals, we detail a regulatory strategy that controls network activity by sensing brain temperature.

## Results

Respiratory-related motor activity can be recorded from the *ex vivo* brainstem-spinal cord via cranial nerve rootlets that innervate muscles of the lower jaw and glottis. Activity from this preparation closely matches the respiratory motor pattern seen in freely-behaving animals (Santin and Hartzler, 2017), allowing the study of the central respiratory network independent from peripheral inputs. As expected, cooling from 22°C to 10°C stopped respiratory motor output (Figure 1A). However, after periods of inactivity between ∼10-20 minutes, 6/9 preparations resumed bursting. After 30 minutes at 10°C, the temperature was warmed to the baseline temperature of 22°C. All preparations experienced an overshoot in burst frequency relative to baseline and then drifted back to control levels by approximately 30 minutes (Figure 1E & F). These responses are consistent with real-time homeostatic regulation of circuit output (Davis, 2013; Ransdell et al., 2012; Styr et al., 2019; Turrigiano et al., 1998): Once cooling stops the network, adjustments in excitability restore activity in most preparations. Upon return to the baseline temperature, compensation during the perturbation manifests as hyperexcitability, which then fades as activity drifts back near the initial value.

**Figure 1.**
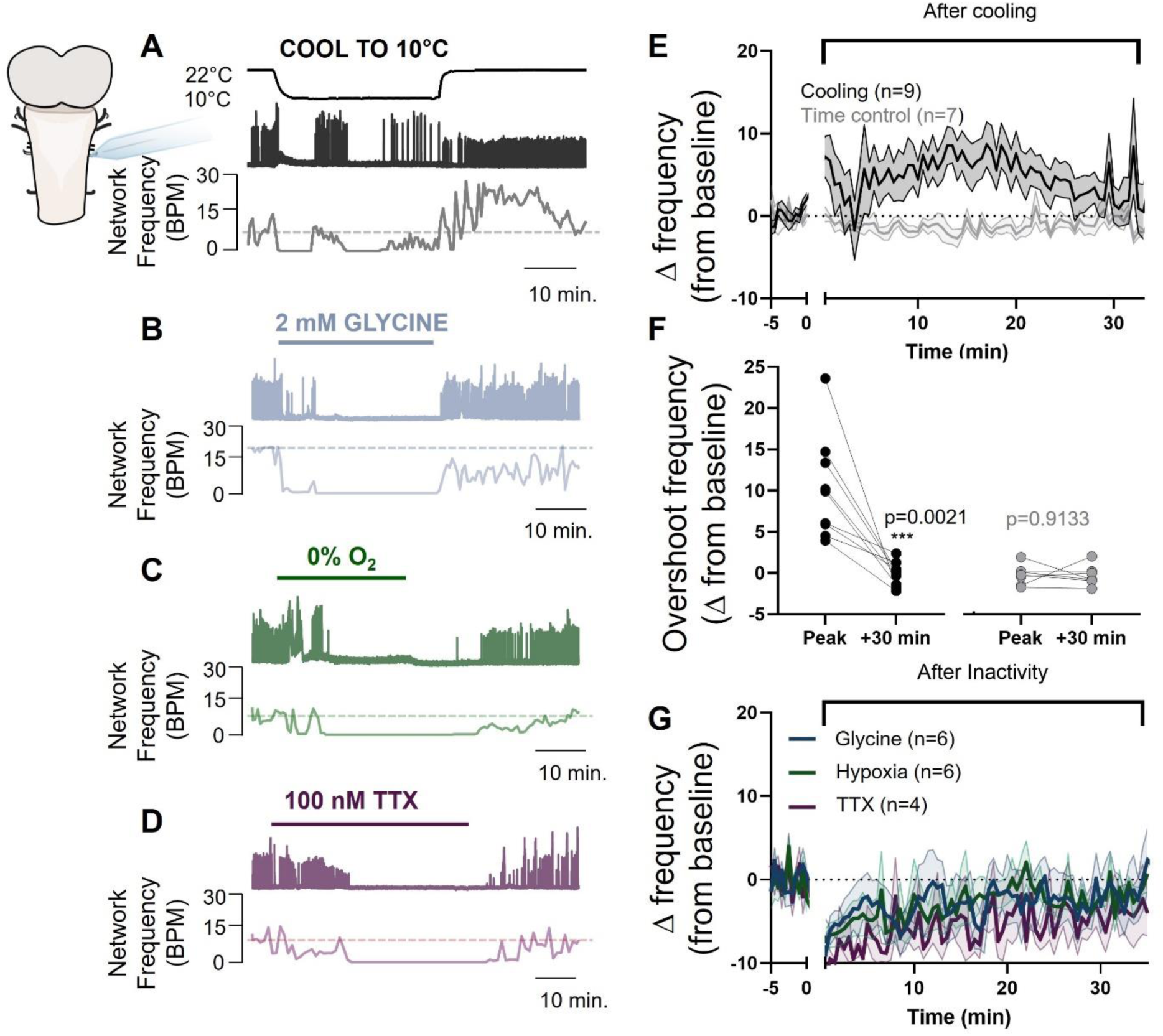
Cold temperature causes inactivity and then induces a homeostatic response that boosts network excitability without sensing activity. A.-D. Example recordings of network inactivity induced by cooling (A). After ∼10 minutes of inactivity by cooling activity recovers. Upon warming the brainstem, activity overshoots the baseline and then recovers in the next tens of minutes. Inactivity treatments at warm temperatures do not show this response (B; 2 mM glycine, C; hypoxia, D; 100 nM TTX). E. Change in instantaneous burst frequency from baseline over the following 30 minutes after cooling (n=9, paired t test), indicating enhanced network excitability compared to baseline following cooling. Time controls left at 22°C do not show this response (n=7, paired t test). F. Averaged data showing the compensatory increase in burst rate following rewarming (left) and no change in time controls (right). G. Summary data from glycine, hypoxia, and TTX, indicating that no compensatory overshoot was observed as seen after cooling.

We hypothesized that the compensatory response following inactivity by cooling may be driven by an activity-dependent signal, consistent with much of the literature in neuronal homeostatic regulation (Wen and Turrigiano, 2024). To test this, we silenced the network at 22°C over the same time course as cooling experiments using 3 approaches: 2 mM glycine to enhance glycinergic synaptic inhibition (n=6), 100 nM tetrodotoxin (TTX; n=4) to block action potentials, and hypoxia (n=6) to stop activity by reducing ATP production, which is likely to occur during the cold. Silencing preparations with glycine and hypoxia did not lead to compensatory bursting (activity could not occur in TTX) (Figure 1B-D). Washing each compound restored activity, but did not lead to an overshoot after activity returned, further indicating that inactivity *per se* does not lead to a compensatory increase in excitability as was consistently observed with inactivity by cooling. Therefore, acute temperature changes, rather than the loss of activity, appear to be the variable that drives compensatory responses.

A key aspect of homeostatic regulation involves Ca^2+^ signaling that acts as a second-messenger (Li et al., 2020). Several ion channels may allow increases in intracellular Ca^+^ during cooling. In amphibians, TRPM8 and TRPV3 are cold-activated channels with Ca^2+^ permeability (Myers et al., 2009; Saito et al., 2011). However, inhibitors of each channel did not alter compensation responses by temperature changes (Supplemental Figure 1A-B&E). In addition, cooling slows the inactivation/desensitization rates of L-type Ca^2+^ channels and NMDA-glutamate receptors by 10-50-fold over 10°C, causing channels to remain open for longer in the cold (Cais et al., 2008; Peloquin et al., 2008). Inhibitors of L-type Ca^2+^ channels and NMDARs failed to oppose the compensation in response to cooling (Supplemental Figure 1C-D&E). Surprisingly, the block of NMDARs led to a greater compensatory overshoot frequency compared to controls, which never recovered to baseline in any experiment (Supplemental Figure 1D&E). In sum, TRPM8, TRPV3, and L-type Ca^2+^ channels are not involved in compensation response, while NMDAR activation appears to function as a constraint.

Neuromodulation is a candidate effector of homeostatic regulation (Khorkova and Golowasch, 2007). The respiratory rhythm of amphibians is thought to be generated by a group of neurons in the reticularis parvocellularis, but is influenced by modulatory systems, including serotonin, norepinephrine, orexin, and more (Milsom et al., 2022). To assess the degree to which compensation involves the core rhythm generating and motor neurons vs. long-range modulatory input, we employed the “thick slice” preparation (Saunders and Santin, 2023). This preparation contains the neurons needed for central pattern generation but lacks much of the neuromodulation present in the intact brainstem. When we cooled thick-slices from 22°C to 10°C and back to 22°C (n=4), we did not observe compensation consistent with that seen in the intact brainstem (Supplemental Fig. 2): Preparations tended to reduce burst frequency upon rewarming, rather than increasing it, indicative of a loss of the compensation response. This suggested that modulatory centers in the intact preparation play a role in enhancing excitability during acute temperature changes.

While many modulatory inputs exist, norepinephrine influences temperature responses of the respiratory network (Vallejo *et al*., 2018). Most neurons have some degree of temperature-sensitivity, typically being slowed by cooling and enhanced by warming in a quasi-predictable manner (Robertson and Money, 2012). However, noradrenergic neurons of locus coeruleus (LC) in amphibians do not follow this trend: Instead, they are strongly *activated by cooling* and *inhibited by warming* via intrinsic mechanisms (Amaral-Silva and Santin, 2023; Santin et al., 2013). Given the paradoxical activation of the LC by cooling, we addressed the role of the midbrain/pontine noradrenergic system as a temperature sensor that boosts excitability of the respiratory network following inactivity by acute cooling.

For this, we first used a chamber that separated pontine/midbrain structures containing the LC from the brainstem (Fig. 2A). Since LC neurons are activated by cold temperatures, we hypothesized that keeping the midbrain/pontine compartment warm would reduce the degree of compensation if activation of these neurons by cooling is involved. When clamping the midbrain/pons at the control temperature and cooling only the brainstem, we observed reduced compensation: Preparations that recovered activity in the cold did so to a lesser extent and did not overshoot baseline frequency upon return to 22°C (Fig. 2A-C). Therefore, neurons within the midbrain/pons must be cooled to fully express homeostatic responses in the brainstem.

**Figure 2.**
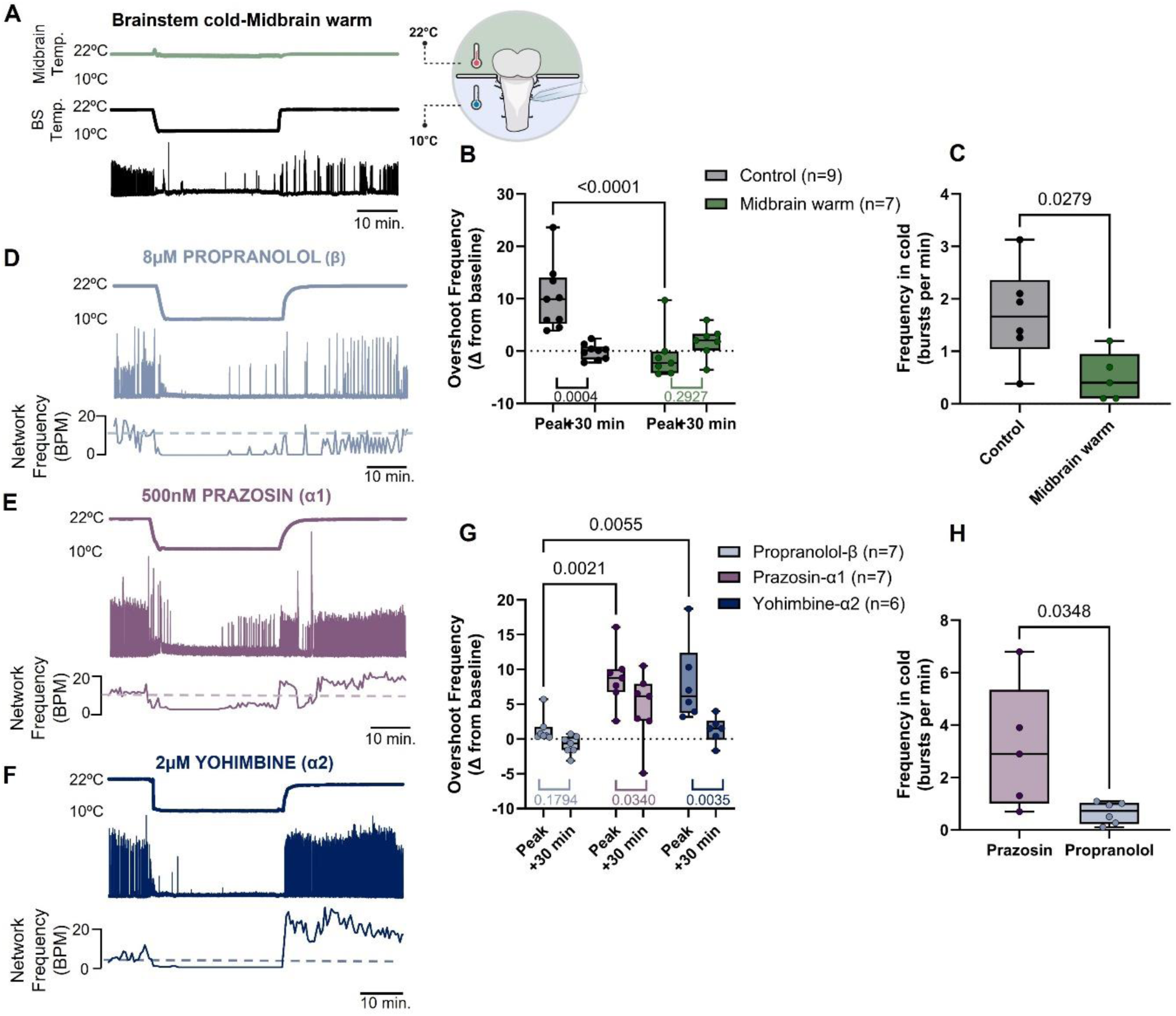
Cooling the midbrain containing noradrenergic neurons, and β-adrenoreceptor signaling is required for homeostatic responses. A. Example recordings showing network responses to acute cooling and warming only the brainstem region while maintaining the midbrain/pons at 22°C. When only cooling the brainstem, no compensatory overshoot response is observed, demonstrating a requirement for midbrain/pontine cooling. D-F show mean data of the peak overshoot response up rewarming (two-way ANOVA followed by Holm-Sidak Multiple Comparisons test) and activity at 10°C (unpaired t test) with cooling only the brainstem (green) compared to cooling the entire brainstem (control; black). B-D. Example recordings of homeostatic responses to acute temperature changes after blocking β, α1, and α2-adrenergic receptors with 8 μM propranolol, 500 nM prazosin, and 2 μM yohimbine. G-H. show mean data of the peak overshoot response up rewarming compared to recovery (two-way ANOVA followed by Holm-Sidak Multiple Comparisons test) and activity at 10°C (unpaired t test) in the presence of each adrenoreceptor inhibitor.

We next determined the contribution of receptors that bind norepinephrine as a cause of compensation. For this, we bath-applied antagonists of α1, α2, and β-adrenergic receptors. α1 and α2 did not play a role, where preparations had typical compensatory responses seen in controls (Fig. 2 E-H). In contrast, the β-adrenoreceptor antagonist, propranolol, reduced the degree of recovery in the cold and strongly blunted the compensatory overshoot upon rewarming (Fig. 2 D, G-H). In addition, activation of βRs using the agonist isoproterenol at warm temperatures mimicked the response to cooling in two ways. First, isoproterenol enhanced network excitability, as demonstrated by an increase in burst frequency. Second, increased excitability had a slow washout time, where burst rate remained elevated after returning to baseline conditions for 30 minutes (Supplemental Fig. 3). Therefore, β-adrenergic signaling is both necessary and sufficient to produce aspects of the network responses observed after acute temperature changes.

Given the requirement for midbrain/pontine cooling and norepinephrine signaling, we addressed how neurons of the LC, which we have recently verified to be noradrenergic (Amaral-Silva and Santin, 2023), transduce cold into enhanced excitability. Corroborating previous results (Santin and Hartzler, 2015), LC neurons in brain slices increase their firing rates during acute cooling (Figure 3A&C), consistent with a requirement for midbrain/pontine cooling and noradrenergic signaling in the compensation response. To determine if these responses were also seen in neurons of the brainstem, we assessed motoneurons from the hypoglossal nucleus. Brainstem motoneurons neurons did not exhibit a firing response to cooling (Supplemental Figure. 4), suggesting specificity of cold-activation within the LC.

**Figure 3.**
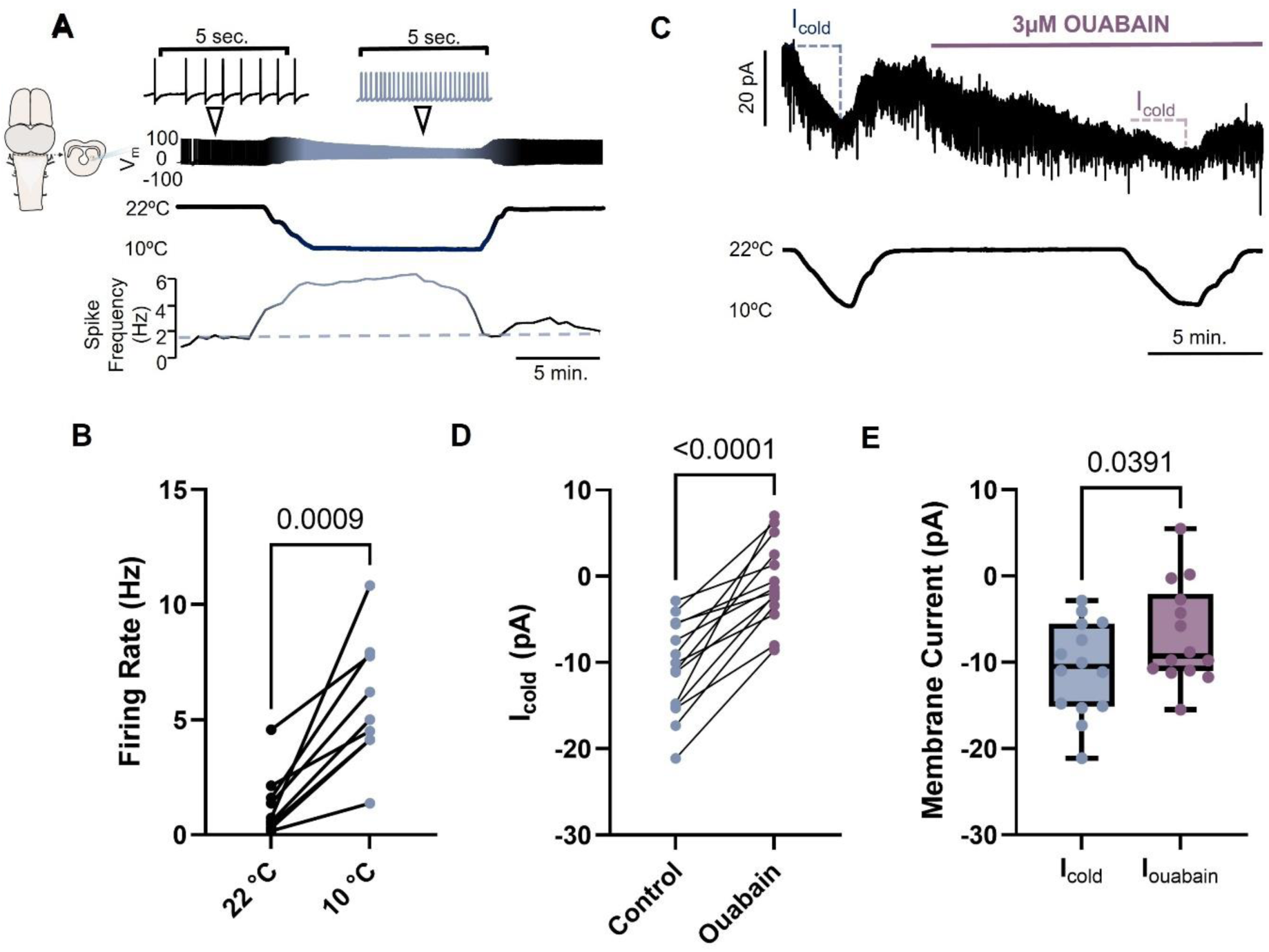
Cold activates neurons within the noradrenergic locus coeruleus *via* inhibition of the Na^+^/K^+^ ATPase. A. Example recording of an LC neuron in a brain slice, demonstrating strong activation by cooling from 22°C to 10°C. B. Mean data showing increases in LC neuron firing rate during cooling (n = 9 cells from N = 4 animals; paired t test). C. Raw voltage clamp recording (v_hold_= - 50 mV) during cooling before and after application of 3 μM ouabain to inhibit the Na^+^/K^+^ ATPase (n = 14 cells from N = 11 animals). Cooling elicits a reversible inward current (I_cold_). Application of ouabain induces an inward current, but cooling in the presence of ouabain reduces the magnitude of I_cold_. D. Mean data showing I_cold_ in aCSF vs. ouabain (paired t test). E. Comparison of I_cold_ vs. I_ouabain_ (paired t test).

For the molecular sensor of cold within LC neurons, we reasoned that TRPM8, TRPV3, NMDARs, and L-type Ca^2+^ channels are unlikely to contribute because these processes did not account for the network homeostatic response. As ATPases are enzymes with potentially high temperature sensitivity (Glitsch, 2001), we focused on how cold may inhibit the tonic hyperpolarizing current from the electrogenic Na^+^/K^+^ ATPase (Na^+^ pump) to stimulate LC neurons. We tested this by voltage clamping the cold-induced current (I_cold_). Cooling LC neurons led to an inward I_cold_, consistent with the ability of cooling to increase depolarizing drive (Figure 3B&D). We then added an inhibitor of the Na^+^ pump, 3 µM ouabain. Ouabain elicited an inward current, indicating inhibition of the tonic outward current from the Na^+^ pump. After reaching steady-state, subsequent cooling in the presence of ouabain produced I_cold_ with a reduced magnitude, and some cells even presented with an outward current instead of an inward current (Figure 3B-D). These results show that inhibition of the Na^+^ pump by cold is necessary to fully express I_cold._ While ouabain also led to an inward current, it was significantly smaller than I_cold_, which suggests that multiple potential mechanisms are at play in generating I_cold_ in addition to inhibition of the Na^+^ pump or 3 μM ouabain inhibits the Na^+^ pump to a lesser extent than 10°C (Figure 3E). Nevertheless, inhibition of the Na^+^ pump serves as a critical transduction step for cooling to stimulate LC neurons, which then elicits network compensation *via* β-adrenergic signaling.

Finally, we sought to link inhibition of the Na^+^ pump to network compensation that occurs during acute temperature changes. We expected that experimentally offsetting inhibition by the Na^+^ pump would reduce compensation. While there are no known selective agonists/activators of the Na^+^ pump, monensin is Na^+^-H^+^ exchanging ionophore, which increases intracellular Na^+^ to activate the Na^+^ pump. This tool has been used in several nervous system preparations, including amphibians (Kueh et al., 2016; Picton et al., 2017; Zhang et al., 2015). To verify this approach, we applied 1 µM monensin to LC neurons in brain slices to determine if activating the Na^+^ pump opposes cold-activation. Application of monensin initially reduced neuronal firing at 22°C, consistent with activation of the outward current carried by the Na^+^ pump. When comparing firing responses to acute cooling in controls, monensin also blunted the cold-activated firing response, suggesting that monensin offsets at least some Na^+^ pump inhibition that stimulates firing (Fig. 4A-C). We then assessed network compensation during experimental activation of the Na^+^ pump by monensin. In response to acute temperature changes, monensin reduced the compensation response: Preparations that recovered activity in the cold did so to a lesser extent and had a less pronounced overshoot upon the return to warm temperatures (Fig. 4D-G). These results show that reversing a key cellular mechanism that causes cold-activation of noradrenergic neurons attenuates the compensatory network response during activity disturbances caused by acute temperature change. In sum, these results point to a feedback loop for the regulation of network activity, where instead of sensing activity, temperature-sensitivity of modulatory neurons leads to compensation within a motor circuit.

**Figure 4.**
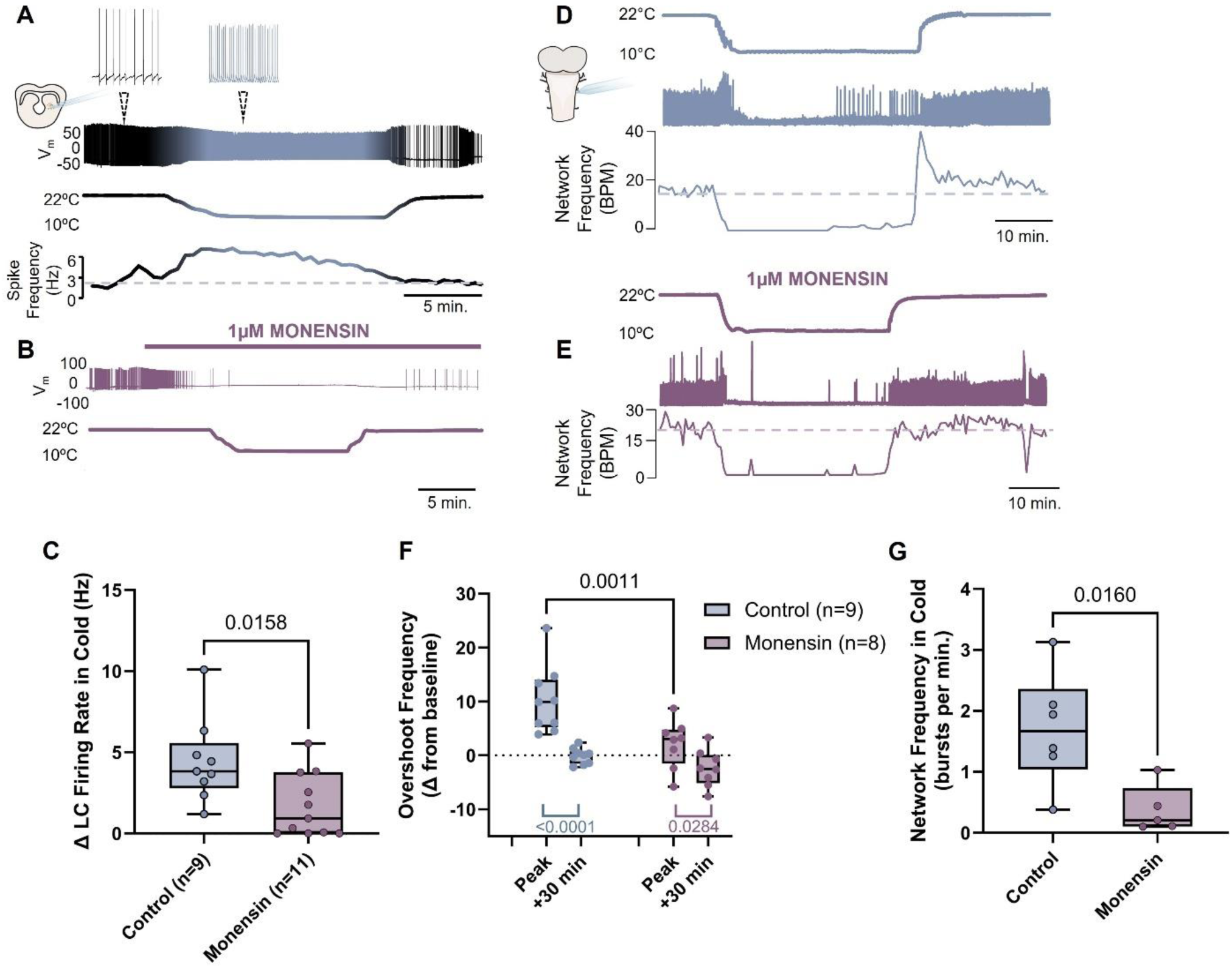
Activating the Na^+^/K^+^ ATPase reduces cold-activated firing of LC neurons and opposes homeostatic recovery of the network. A-B. Example recordings of LC neurons in aCSF (control) and in the presence of 1 μM monensin to activate the Na^+^/K^+^ ATPase (n = 11 cells from N = 6 animals). Control neurons strongly increase firing rate. With monensin, activity slows, consistent with activating of the Na^+^/K^+^ ATPase, and stimulatory response to cooling is blunted. C. Mean data comparing the cold-induced firing responses of LC neurons in aCSF and monensin (unpaired t test). D-E. Example recordings of homeostatic responses during acute temperature changes in aCSF (control) and in the presence of 1 μM monensin. F-G show mean data of the peak overshoot response up rewarming compared to recovery (two-way ANOVA followed by Holm-Sidak Multiple Comparisons test) and activity at 10°C (unpaired t test) in controls and 1 μM monensin.

## Discussion

Over 30 years ago, an important set of studies suggested that neurons transduce their own activity levels, in turn, causing cellular adjustments aimed at controlling network output through activity-dependent homeostasis (LeMasson et al., 1993; Turrigiano et al., 1994). This set in motion a blueprint for explaining how the healthy nervous system maintains stable activity patterns, and how issues with these regulatory systems could disrupt circuit output in disease (Chen et al., 2022). A core interpretation of nearly all models of feedback homeostasis in the nervous system is that some aspect of neural activity—be it intracellular Ca^2+^ dynamics, neurotransmitter receptor activation state, or membrane potential— is vital for regulation (Fong *et al*., 2015; O’Leary et al., 2014; Santin and Schulz, 2019). Here, our results diverge from this theme and introduce that specific environmental cues, in this case temperature, may trigger homeostatic responses without tracking activity of the circuit.

To achieve regulation in this way, our results demonstrate an important role for modulatory neurons as temperature sensors (Fig. 2A), β-adrenoreceptor signaling (Fig. 2D, G-H), and a built-in constraint by NMDA-glutamate receptors (Supplemental Fig. 2D-E). The respiratory rhythm in adult amphibians is thought to be generated by mechanisms that involve reciprocal inhibition, as well as excitatory neurons that carry this rhythm to motor pools (Kottick et al., 2013; Saunders and Santin, 2024). Therefore, excitatory metabotropic signaling from β-adrenoreceptors and constraint by NMDARs likely acts somewhere along this path. To activate β-adrenoreceptors in a way that enhances excitability in the brainstem, we show a key step is midbrain/pontine cooling, which seems to involve the activation of noradrenergic neurons via inhibition of the Na^+^ pump (Fig. 3-4). As Na^+^ pumps are important transporters in all neurons, and brainstem motoneurons do not exhibit such strong activation by cooling (Supplemental Fig. 5), we suggest that features of the Na^+^ pump may be unique within the LC. The α subunit of the Na^+^ pump undergoes substantial RNA editing, which gives rise to a range of functionally diverse proteins (Palladino et al., 2003). Therefore, we speculate that RNA editing of the Na^+^ pump within LC neurons may contribute to cold-sensitivity, in addition to interactions with features of LC neurons to enhance firing in the cold [e.g., high input resistance (Santin and Hartzler, 2015)]. Regardless of implementation of these specific mechanisms, or other potential mechanisms, temperature sensing within modulatory neurons defends homeostasis of the brainstem motor network.

Why use an environmental variable like temperature to regulate the activity of a motor circuit? We suggest that this strategy may build specificity for regulation in different environments and allow circuits to implement regulation across timescales. For specificity, while breathing is a critical rhythmic behavior, amphibians use peripheral sensory systems to inhibit ventilation, potentially, as a predator detection and avoidance strategy (Coates and Ballam, 1990; Santin and Hartzler, 2016). If sensors of global network activity induced compensation over the timeline we observed here, the homeostatic response that followed could be detrimental to the goal of predator avoidance by haphazardly restarting or increasing breathing. In this way, sensitivity to specific environmental variables seems to allow the network to restore activity in conditions where it is needed (cool temperatures), while leaving the ability to keep the network silent in other situations where it is adaptive to do so.

Using different signals, such as temperature, may also allow circuits to perform regulation across different timescales. A problem in neuronal regulation has been identified, where it is difficult to maintain homeostasis with multiple sensors operating over different time intervals that use the same activity signal (e.g., altered intracellular Ca^2+^) (Liu et al., 1998; O’Leary *et al*., 2014) [but see a recent study that offers a theoretical solution to this problem (Alonso *et al*., 2023)]. Results in the amphibian respiratory network have begun to accumulate, indicating that separating regulation mechanisms by induction cue may offer a workaround to this problem. On short timescales (seconds to minute), respiratory motoneurons sense increases in activity through rises in intracellular Na^+^ and membrane depolarization via activation of an isoform of the Na^+^ pump with low Na^+^ affinity and Kv7 channels, respectively (Pellizari et al., 2023). During inactivity over weeks to months seen in hibernation, synaptic compensation also enhances motoneuron and network excitability using both inactivity and Ca^2+^-dependent signals (Zubov *et al*., 2022). Temperature sensing at modulatory neurons which activates metabotropic receptors as we show here may provide a way to restrict regulation to intermediate time scales (tens of minutes) without interfering with faster and slower processes. Therefore, using different induction cues for regulation seems to endow circuits with the capacity to meet different regulatory challenges of varying duration and type.

These results point to organizational rules for regulation that may be applicable to mammals and beyond. For example, several circuits have been reported to lack compensation *via* activity-dependent signaling, at least over certain time scales (Begum et al., 2016; Hamood et al., 2015; Yu et al., 2006). It is possible that regulation may be achieved instead by transducing select variables from the neuronal microenvironment. While the temperature perturbation used here is relevant to amphibians and other poikilothermic species (*i.e*., all animal groups expect for most mammals and birds), certain mammalian brain regions such as the hippocampus and nucleus accumbens can vary by as much as 4°C Celsius depending on the level of metabolic activity (Kiyatkin, 2011), in addition to other microenvironment shifts such as pH and extracellular K^+^ (Chesler, 2003; Moody Jr et al., 1974). Beyond environmental sensors for feedback control, these processes likely work in tandem with network designs that build resilience to environmental perturbations to prevent activity disturbances before they occur (Saunders and Santin, 2024; Tang et al., 2010; Tang et al., 2012). Overall, new insights into how neural circuits accomplish the feat of maintaining stability despite ever-changing internal and external environments will likely be found along a path that integrates activity-dependent, as well as environmentally-driven feedback processes.

## Acknowledgements

This work was funded by the National Institutes of Health (R01NS114514 to JS). We would like to thank Thamar Scipio and Natalie Heath for performing the isoproterenol and yohimbine experiments, respectively. We also would like to thank Nik Bueschke for technical assistance throughout the project and training on patch clamp electrophysiology.

## Materials & Methods

### Animals

All protocols were approved by the Animal Care and Use Committee (ACUC) at the University of Missouri (protocol # 39-264). Adult female American Bullfrogs (n=119), *Lithobates catesbeianus,* purchased from Rana Ranch (Twin Falls, Idaho), were used for all experiments. Frogs were housed in 20-gallon plastic tanks filled with dechlorinated tap water that was perpetually bubbled with room air, with access to both wet and dry areas. Water was cleaned daily and changed as needed. Frogs were fed pellets once a week and were kept on a 12hr:12hr light:dark cycle, with all experiments taking place during the light cycle.

### Drugs

Several pharmacological tools were used in this study: 2 mM Glycine (dissolved in DI water), 10 μM verapamil (dissolved in DI water), 20 μM D-APV (dissolved in DI water), 3 μM ouabain (dissolved in ethanol), 2 μM monensin (dissolved in ethanol), 2 μM Tripvicin (dissolved in DMSO), 100 nM TTX (diluted from 10mM; dissolved in DI water), and 1μM (-)-Bicuculline methiodide (dissolved in DI water) were from Hello Bio (Princeton, NJ, USA). 8 μM propranolol (diluted from 50mM; dissolved in DI water), 500 nM prazosin (dissolved in DI water), and 2 μM yohimbine (dissolved in DI water) were from TCI America (Portland, OR, USA). 3 μM strychnine hydrochloride (dissolved in DI water) was from Sigma-Aldrich (St Louis, MO, USA). 500 nM AMG 333 (dissolved in ethanol) was from Tocris.

### Brainstem-spinal cord preparation

Frogs were deeply anesthetized using approximately 1 ml of isoflurane in a closed 1 L chamber until response to toe pinch was no longer observed. Rapid decapitation with a guillotine was utilized to achieve euthanasia. Brainstem dissections were performed in ice-cold artificial cerebrospinal fluid (aCSF). The forebrain was pithed immediately, and the brainstem-spinal cord was carefully extracted while preserving the nerve roots. The remains of the forebrain were then separated, and the brainstem was cut caudal to spinal nerve III. Following the dissection, the brainstem was pinned to a ∼6mL Sylgard-coated dish and superfused with oxygenated aCSF using a peristaltic pump. In a subset of experiments, the brainstem was pinned in a “split bath” dish containing walls protruding from either side, creating two separate chambers superfused by two separate pumps with two separate temperature controls. In this dish, the brainstem was positioned between the two walls, with the midbrain/pontine region sitting on one side, and the brainstem on the other. An agar block was placed between the two walls and positioned above the brainstem preparation close enough to effectively modulate each chamber’s temperature, but far enough to allow sufficient perfusion to that region. For extracellular nerve recordings, near-synchronous respiratory-related activity can be observed on cranial nerves V, VII, X, and hypoglossal. Either cranial nerve X (vagus) or hypoglossal activity was recorded using a glass suction-electrode, and AC amplified (1000x, A-M Systems Model 1700, A-M Systems, Carlsborg, WA, USA), filtered (10Hz-5kHz), and digitized (PowerLab 8/35 ADInstruments, Colorado Springs, CO, USA).

### Experimental protocols

Nerve output was allowed to stabilize for approximately 1 hour at room temperature before drug application or temperature change. In control experiments (n=9), the temperature of the aCSF was decreased from 22°C to 10°C using a bipolar in-line temperature controller (Warner Instruments Model CL-100, Warner Instruments, Hamden, CT, USA), and the temperature of the solution was monitored with a thermocouple positioned close to the brainstem. The temperature was held at 10°C for 30 minutes before warming back up to the baseline temperature of 22°C.

A subset of experiments was performed to induce network inactivity in control temperatures. In one group (n=6), 2mM glycine was added to the perfusate and allowed to superfuse for 30 minutes before getting washed out. A different group (n=4) was exposed to 100nM tetrodotoxin for ∼60 min prior to washing out. TTX was used for 60 minutes because 100 nM took ∼20-30 minutes to fully stop activity, which allowed us to more closely match the duration of inactivity near 30 minutes. A final group (n=6) was exposed to anoxia for 25 min prior to washing out. We selected 25 minutes because most preparations took ∼5 minutes to restart upon the return of O_2_, which again allowed us to match the duration of inactivity. Hypoxia was achieved via superfusion of aCSF that was bubbled with a gas mixture that contained 0% oxygen, 98.5% nitrogen, and 1.5% carbon dioxide. The temperature was kept at 22°C for the duration of each of these experiments. For all experiments that used various inhibitors, each was applied for 10-15 minutes before cooling and then followed the normal protocol to cool and rewarm the tissue.

### Thick slice preparation

To address the capacity for a minimal unit of the respiratory network to undergo the compensation response, we used a recently developed thick-slice preparation (Saunders and Santin, 2023). This preparation exhibits features of the intact brainstem preparation (burst pattern and stereotyped sensitivity to modulators as seen in the intact brainstem such as suppression by opioid agonist and stimulation by serotonin) but produces activity at ∼1/10 of the normal rate. Following dissection of the intact brainstem, vagus nerve output from the intact brainstem was briefly observed preceding transection to ensure preparations were viable. Using small spring scissors, the brainstem was transected rostral and caudal to the vagus nerve root. Shortly following transection, 3µM strychnine + 1µM bicuculine was added to the aCSF perfusate which is needed to trigger burst activity (Saunders and Santin, 2023). Vagal motor output was given >30 min to stabilize prior to decreasing the temperature from 22°C to 10°C. Following a 30-minute period at 10°C, the temperature was increased back to baseline and activity was recorded for another 30-40 minutes.

### Whole-cell patch clamp electrophysiology

Following dissection, brainstem preparations were adhered to an agar block mounted on a vibratome tissue slicer and submerged in cold, bubbled aCSF. 300 µM transverse slices were obtained from regions containing either the locus coeruleus (LC) or hypoglossal motor nucleus and left to recover in room-temperature, bubbled with aCSF. Upon transfer to the recording chamber, slices were immobilized with a nylon grid and kept with constant oxygenated aCSF super-perfusion. Neurons were visualized with a fixed-stage microscope (FN1, Nikon Instruments Inc., Melville, NY, USA). Glass patch pipettes were pulled using a P1000 horizontal pipette puller (Sutter Instruments, Novato, CA, USA) and backfilled with artificial-intracellular solution composed of the following (in mM): 110 K-gluconate, 2 MgCl_2_, 10 HEPES, 1 Na_2_-ATP, 0.1 Na_2_-GTP, 2.5 EGTA. Pipettes were positioned near the neurons of interest using a micromanipulator (Sutter Instruments, Novato, CA, USA) and whole-cell recordings were acquired using an Axopatch 200B amplifier and Axon Digidata 1550 digitizer (Molecular Devices, San Jose, CA, USA).

### LC neuron experiments

Neurons found in the area thought to be homologous to the locus coeruleus of mammals (González et al., 1994; Marin et al., 1996), which also express dopamine beta hydroxylase for norepinephrine synthesis (Amaral-Silva and Santin, 2023), have previously been found to increase firing rates during acute cooling (Santin *et al*., 2013). To establish the response of LC neurons to cold temperatures, a subset of cells (n=9) was recorded in current-clamp mode (I = 0) and allowed to establish a baseline firing-rate (∼5 min.). In cells that did not spontaneously fire, a small positive current was introduced and increased until the cell fired and remained firing at a consistent rate. Once baseline was established, the temperature of the chamber was decreased from 22°C to close to 10°C (temperature varied 1-2° between cells depending on the probe’s position in the chamber) using a bipolar in-line temperature controller (Warner Instruments). Temperature remained around 10°C for 10 minutes before the chamber was brought back to the baseline temperature.

In a different set of experiments, LC neurons (n=14) were recorded in voltage-clamp mode (holding voltage = -50mV) and allowed to establish a baseline current. After baseline was established, the temperature of the chamber was decreased from 22°C to around 10°C for a brief period prior to returning to the baseline temperature. The cell was given several minutes to stabilize at 22°C before the slice was exposed to 3µM ouabain in aCSF. Ouabain was continuously supplied into the chamber for long enough to observe an inward current in most cells. When this inward current saturated, a second cooling was performed in the presence of ouabain.

In a separate subset of current-clamp recordings, LC neurons (n=11) were exposed to 1µM monensin following an initial stabilization period (>5 min.). After several minutes (>5 min.) of monensin exposure, the temperature of the chamber was decreased from 22°C to 10°C, where it remained for a duration of 10 min. and then returned to baseline temperature for an additional 10-minute period.

### Hypoglossal motoneuron experiments

A series of hypoglossal motoneuron current-clamp recordings (n=9) were obtained to compare to the temperature response of cells in the locus coeruleus. Neurons were given time to establish a baseline firing rate (5 min.) prior to being cooled from 22°C to 10°C for a 30-minute period. The temperature was then brought up to baseline for an additional 15 minutes.

### Data Analysis

#### Brainstem preparations

Burst frequency for 5 minutes at baseline, in the last 10 minutes of cold, the peak 5 minutes during the overshoot, and then burst frequency at 30 minutes were analyzed for all network level experiments. In experiments that used various pharmacological tools, these were applied for 10-15 minutes before cooling. To quantify the overshoot we analyzed the change in burst frequency compared to baseline. While the starting frequency across preparations differed, the change from baseline during the overshoot period did not correlate with starting frequency (i.e., we observed similar absolute changes regardless of the baseline activity level) (Supplemental Fig. 5). Thus, we performed analyses on the absolute change from baseline. While nearly all preparations exhibited an overshoot following cooling suggestive of compensation in the cold (e.g., all groups except monensin and propranolol), a minority of these preparations did not show activity during cold temperatures. Therefore, when comparing burst activity in the cold under different conditions, we only used preparations that recovered activity. If groups did not have at least n=5 preparations that recovered in the data set, no analysis during the cold was performed.

#### Whole-cell patch clamp

For control LC current-clamp experiments, we analyzed the spike frequency for 30 seconds during the baseline period and the period in the cold where frequency was at its peak. The same was done in current-clamp experiments in which LC neurons were exposed to monensin with the addition of a 30 second period after monensin exposure, but before the temperature was reduced.

In voltage-clamp experiments, the change in membrane current between baseline and the first period of cold (I_cold_) was analyzed. Then, after an inward current produced by ouabain was observed, the change in membrane current induced by ouabain (between the recovery and immediately prior to the second period of cooling) was taken as “I_ouabain_”.

### Statistics

Activity during the cold was compared with an unpaired t test. To compare changes in burst activity during the overshoot and recovery to baseline, we used repeated measures two-way ANOVA and Holm-Sidak Multiple Comparisons tests. For comparisons of only one group with multiple time points (isoproterenol experiments), we used a one-way repeated measure ANOVA, followed by Holm-Sidak Multiple Comparisons tests. Firing rate data before and after cooling, and I_cold_ before and after ouabain, were compared using a paired t test. Comparisons of I_cold_ vs. I_ouabain_ and LC, hypoglossal response to cooling, the change in firing in control vs. monensin used unpaired t tests. Significance was accepted with p<0.05.

## Supplementary Figures

**Supplementary Figure 1.**
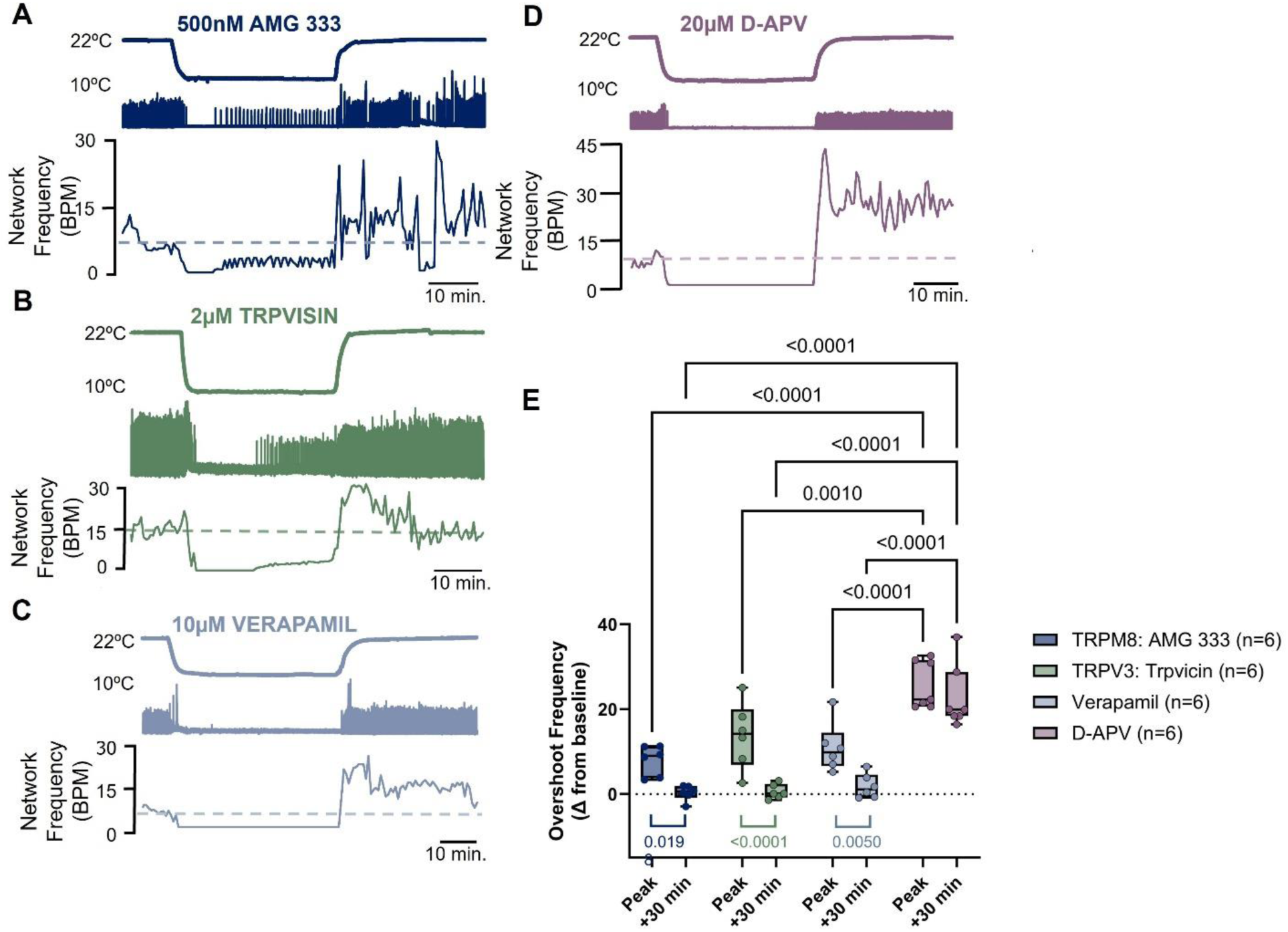
ThermoTRP channels and L-type Ca^2+^ channels do not contribute to homeostatic responses, while NMDARs constrains them. A-D shows raw recordings of the cooling protocol in the presence of AMG 333, trpvicin, verapamil, and D-AP5 to block TRPM8, TRPV3, L-type Ca^2+^ channels, and NMDARs. AMG 333, trpvisin, and verapamil showed similar qualitative responses as a control preparations, with an overshoot average cooling, which trends back to baseline over the next 30 minutes. D-APV, however, led to strongly exaggerated overshoots that did not recover to baseline. E. Mean data of change in burst frequency compared to baseline (overshoot) and recovery to baseline (two-way ANOVA followed by Holm-Sidak multiple comparisons tests).

**Supplementary Figure 2.**
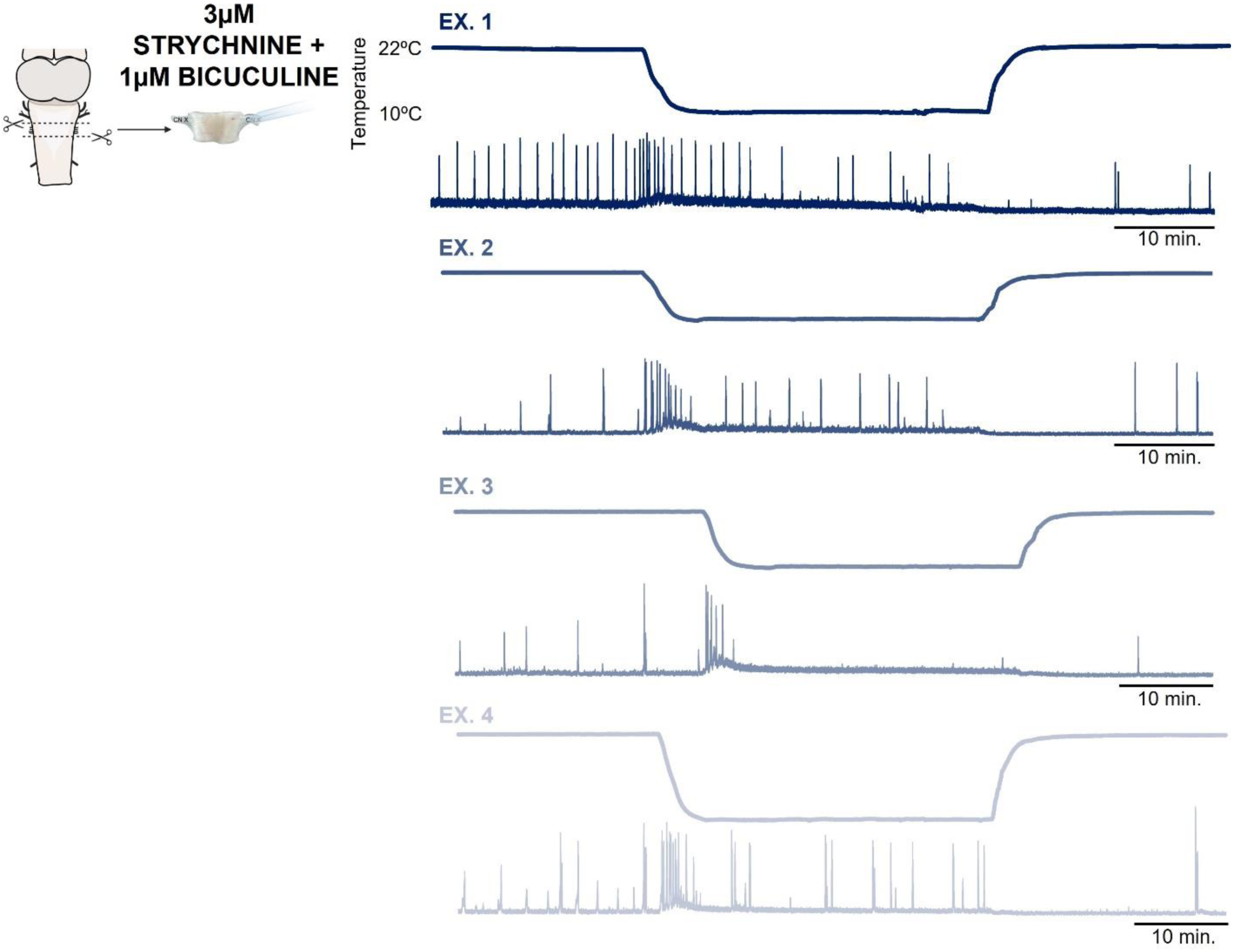
A section of brainstem that lacks long-range modulatory centers does not elicit compensation after cooling. Raw recordings of 4 “thick slice” preparations that produce respiratory-like activity. During a 30 minute cooling protocol, burst activity was variability influenced by temperature, where 1 preparation stopped and 3 did not. However, none exhibited a compensatory overshoot during the 30 minutes recovery period as commonly observed in the intact brainstem preparation. This suggests that modulatory centers are needed to observe the typical responses to temperature in the intact brainstem.

**Supplementary Figure 3.**
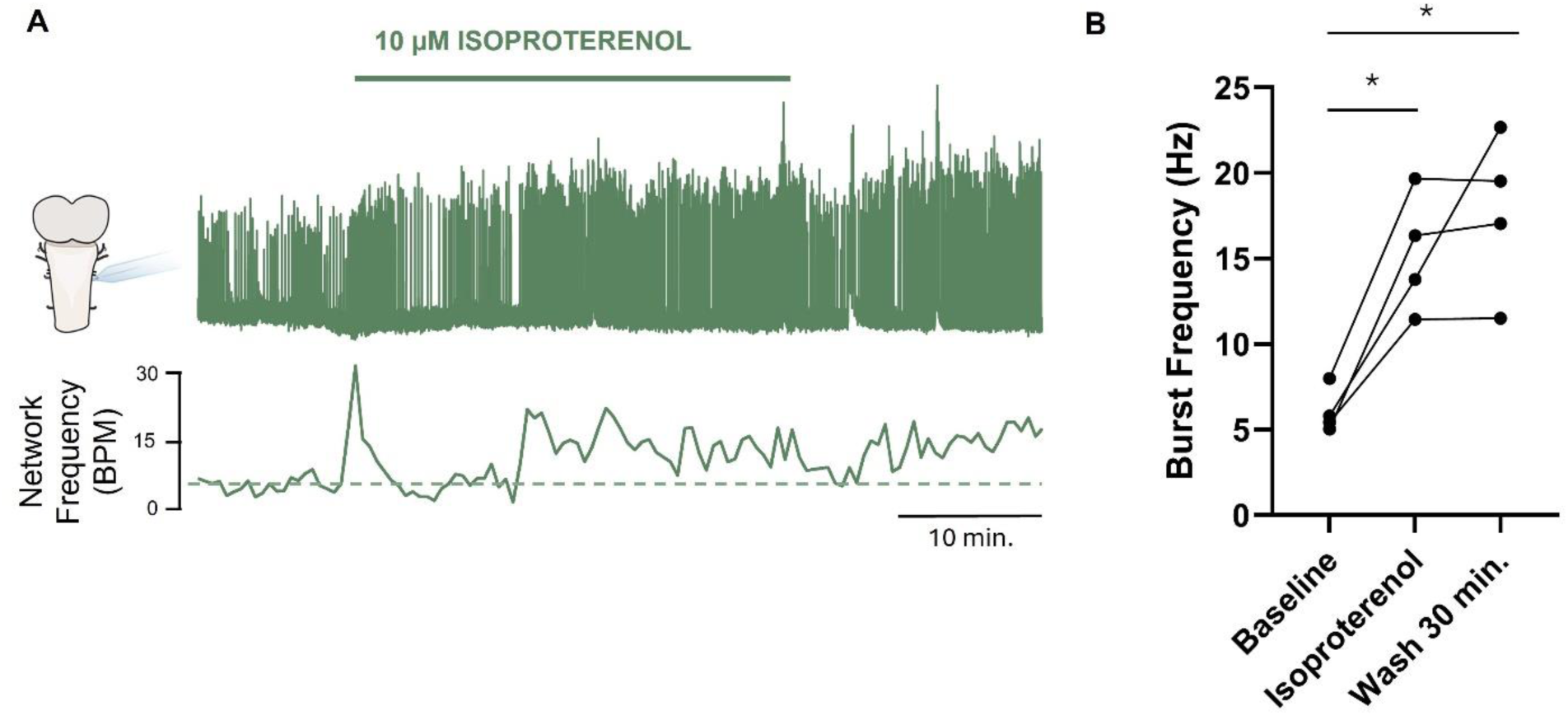
Activating β-adrenoreceptors leads to a prolonged enhanced of network activity. 10 μM isoproterenol stimulates respiratory network frequency and displays slow washout kinetics. Therefore, activation of β-adrenoreceptors is sufficient to explain increases in network excitability that persist and fade slows after beyond the removal of cold stimulus. N=4 experiments, one-way ANOVA followed by Holm-Sidak Multiple Comparisons test. * indicates p<0.05.

**Supplementary Figure 4.**
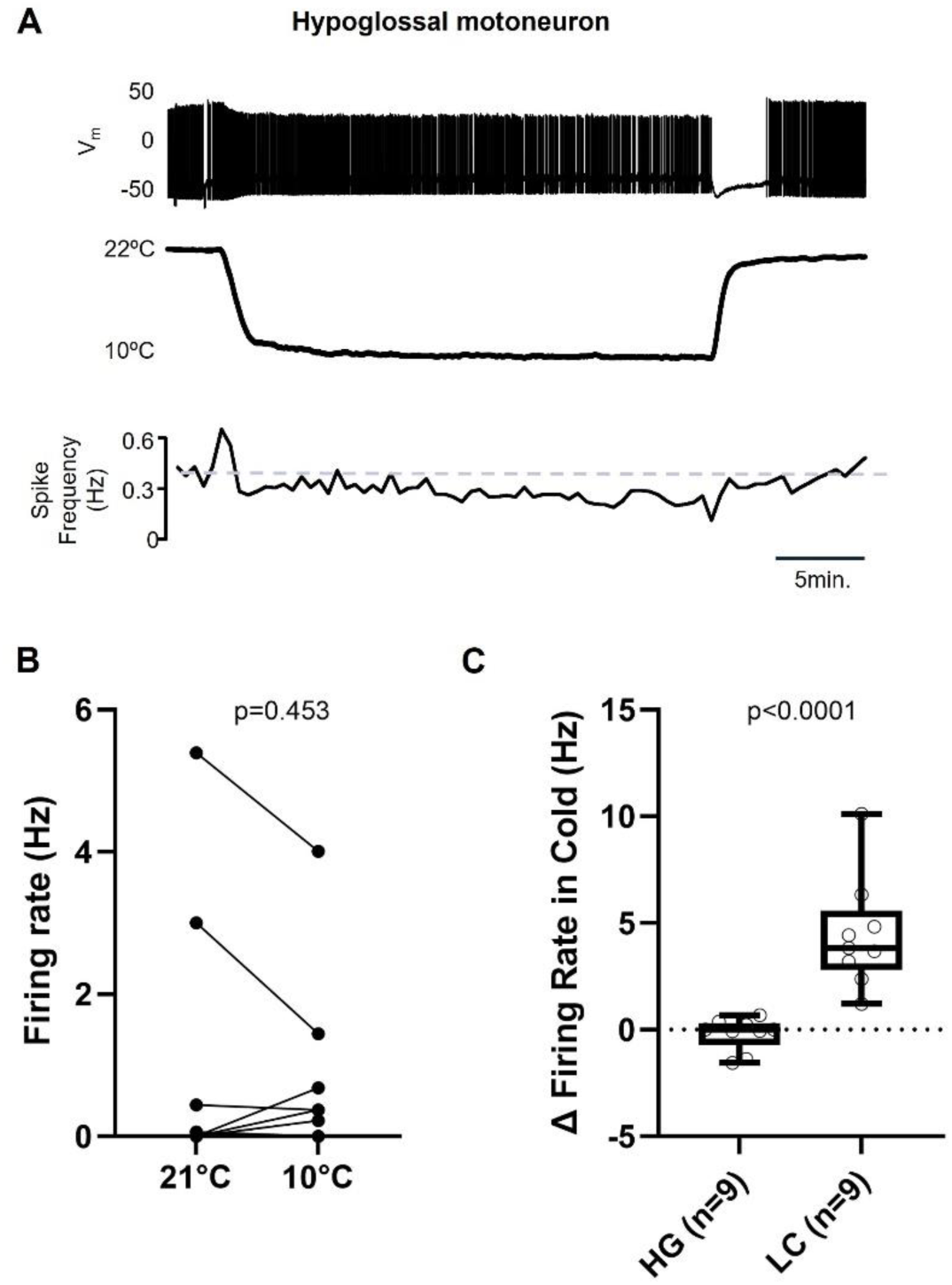
Brainstem motoneurons are not activated by cooling like LC neurons. A. Example recording of a hypoglossal motoneuron in response to cooling. Unlike the LC, motoneurons do not increase firing rates in the cold. B. Mean data of n=9 neurons, showing that firing rates of hypoglossal motoneurons are not significantly increased in response to cooling. C. A direct comparison of LC neurons vs hypoglossal motoneurons indicates LC neurons have enhanced firing rates in the cold compared to brainstem motoneurons (unpaired t test).

**Supplementary Figure 5.**
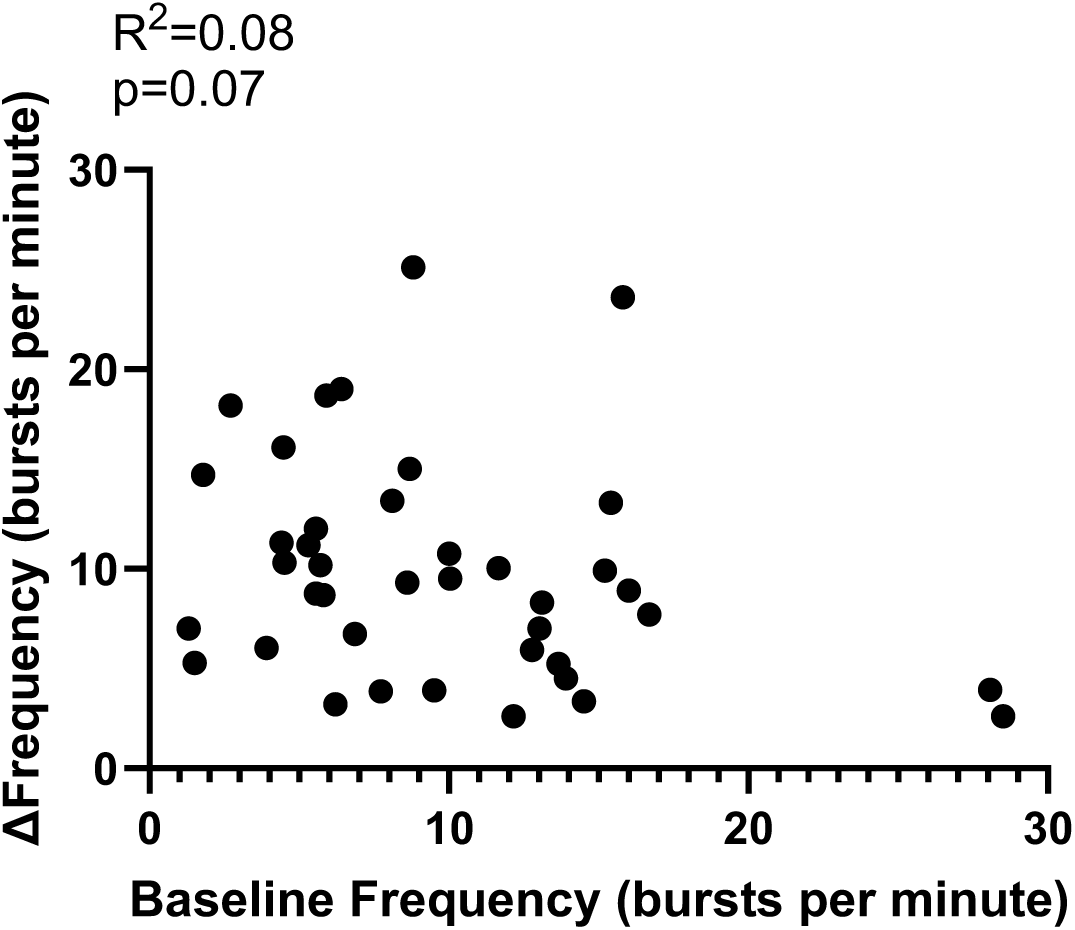
The change in frequency compared to baseline in the overshoot phase does not correlate with baseline frequency.

## Notes

### Competing Interest Statement

The authors have declared no competing interest.

